# Fine scale prediction of ecological community composition using a two-step sequential machine learning ensemble

**DOI:** 10.1101/2021.03.24.436771

**Authors:** Icíar Civantos-Gómez, Javier García-Algarra, David García-Callejas, Javier Galeano, Oscar Godoy, Ignasi Bartomeus

## Abstract

Prediction is one the last frontiers in ecology. Indeed, predicting fine scale species composition in natural systems is a complex challenge as multiple abiotic and biotic processes operate simultaneously to determine local species abundances. On the one hand, species intrinsic performance and their tolerance limits to different abiotic pressures modulate species abundances. On the other hand there is growing recognition that species interactions play an equally important role in limiting or promoting such abundances within ecological communities. Here, we present a joint effort between ecologists and data scientists to use data-driven models informed by ecological deterministic processes to predict species abundances using reasonably easy to obtain data. To overcome the classical procedure in ecology of parameterizing complex population models of multiple species interactions and poor predictive power, we followed instead a sequential data-driven modeling approach. We use this framework to predict species abundances over 5 years in a highly diverse annual plant community. Our models show a surprisingly high spatial predictive accuracy (RSE ~ 0.13) using only easy to measure variables in the field, yet such predictive power is lost when temporal dynamics are taken into account. This result suggest that predicting the temporal dimension of our system requires longer time series data. Such data would likely capture additional sources of variability that determine temporal patterns of species abundances. In addition, we show that these data-driven models can also inform back mechanistic models of important missing variables that affect species performance such as particular soil conditions (e.g. carbonate availability in our case). Being able to gain predictive power at fine-scale species composition while maintaining a mechanistic understanding of the underlying processes can be a pivotal tool for conservation, specially given the human induced rapid environmental changes we are experiencing. Here, we document how this objective can be achieved by promoting the interplay between classic modelling approaches in ecology and recently developed data-driven models.

**Author summary:** Prediction is challenging but recently developed machine learning techniques allow to dramatically improve prediction accuracy in several domains. However, these tools are often of little application in ecology due to the complexity of gathering information on the needed explanatory variables, which often comprise not only physical variables such as temperature or soil nutrients, but also information about the complex network of species interactions regulating species abundances. Here we present a two-step sequential modelling framework that overcomes these constraints. We first infer potential species abundances training models just with easily obtained abiotic variables, and then use this outcome to fine-tune the prediction of the realized species abundances when taking into account the rest of the predicted species in the community. Overall, our results show a promising way forward for fine scale prediction in ecology.

## Introduction

In the face of human-induced rapid environmental change, the ability to predict species responses to different drivers of environmental change within a community context is more pressing than ever [1]. However, fine scale prediction is a recognized weak spot in ecology [2–5]. Within the realm of community ecology, most prediction efforts rely on a mechanistic understanding of how multiple abiotic and biotic processes regulate species population dynamics. In particular, theoretical frameworks centered around the study of the determinants of species coexistence have experienced during the last decades a fast growth due to the development of concepts and associated mechanistic models investigating the effect of the environment and species interactions on the maintenance of biodiversity [6]. These recent developments have been pivotal to point out ecological processes regulating the dynamics of interacting species such as those occurring in plant competitive networks [7–9]. Moreover, this body of theory has also shown direct applications to better predict species abundances under controlled experimental conditions [10, 11]. Yet, current theory and associated modelling tools fail in most cases to accurately predict basic features of ecological communities observed in nature such as species abundances, composition, and species turnover in space and time [12]. In order to solve this limitation, there is a recent call to address the complexity of multispecies processes occurring in nature [13, 14]. However, a major stumbling block to advance in this front is parameterizing and validating those models in real communities, which currently is prohibitive due to the complexity of estimating with confidence all parameters from observational data [15]. To solve this trade-off between model complexity and data availability, we aim to develop an alternative approximation using a mechanistically informed data-driven approach that allow us to achieve predictive power with affordable data requirements.

In a nutshell, existing phenomenological approaches that summarize well-known mechanistic processes require to feed models describing the population dynamics of interacting species with information about 1) the intrinsic ability of species to grow in the absence of interactions, 2) the strength on intra- and inter-specific interactions, and 3) how these two sets of parameters change in the presence of different abiotic and biotic variables such as soil conditions or multitrophic species interactions (e.g. pollinators, herbivores) [16, 17]. This is in most cases unfeasible for two reasons: 1) we need to gather detailed information under natural conditions, which for many systems is unfeasible due to the long lifespan of species or the inability to detect and quantify the strength of species interactions, and 2) this approach considers that all species within a community can potentially interact among them [15, 18]. As the number of parameters to estimate scales exponentially with the number of species in the community, estimating all parameters for large communities quickly becomes an intractable problem. Moreover, because species abundances are not likely to vary independently (i.e. the population size of species A, B and C covary), it is often difficult to estimate with confidence the strength and sign of many interspecific parameters. Even if we find a suitable ecosystem to parameterize these models, gathering all required information is labor intensive and highly time consuming. Hence, to resolve this conundrum, we can not rely simply in gathering more and better data. We also need simpler models and search for indirect methods to obtain enough information to be predictive. A key challenge, for example, is that the high level of abstraction of mechanistic models do not always translate well to the empirical data we can easily measure [19]. Hence, we need models that move closer to what we could actually measure on the field. But how to capture complex systems with simpler models?

Fortunately, there is a possibility worth exploring. The problem of inferring key behaviours from complex data has been solved using machine learning approaches. Machine learning is a field of computer science that gives computers the ability to learn without being explicitly programmed. In the past decade, machine learning has given us self-driving cars, practical speech recognition, effective web search, and a vastly improved understanding of the human genome [20–23]. However its potential has been unleashed mostly in applied domains, as predictions done by machine learning often lack the interpretability needed to explain the mechanisms behind the algorithm’s decisions. As scientists, we are often uncomfortable with predictions that have no theoretical basis. This implies that we need to combine the power of data-driven models with a stronger interpretability [24]. Here we address this issue by partnering together ecologists and data scientists to develop an efficient and predictive data-driven model rooted in known ecological mechanisms that are thought to explain species occurrence and its abundance at local scales. First, we explain the core problem, then we propose a solution, and finally, we test the predictions against a well-resolved dataset consisting on 5 years of observations describing the community composition of 23 species co-occurring in a Mediterranean annual grassland.

## The problem

To predict species abundances within a community context, we know that different abiotic factors determine species performance and their tolerance limits [25], from which one can derive potential species abundances [26]. However, we also know that the final species fate will be modulated by the positive and negative species interactions established among and within species able to grow in a particular place [27, 28]. Of course, stochastic process coming for instance from dispersal events or random birth and death [29, 30] are also recognized to have an increasing importance in modulating species persistence, but for a first approximation and for the sake of simplicity they are not included into the modelling approach here developed. Hence, mechanistic models to understand species population dynamics and their ability to persist in the long-run are often formalized as a set of coupled equations where each response variable (i.e. population size of a given species in a given time and location) depends on and modifies the outcome of the rest of response variables (i.e. population size of this and other species) [27, 28]. A clear example using the standard Lotka-Volterra equations is the persistence of the populations of three plant species following rock-scissors-paper dynamics [31, 32], in which each species have to win and to lose simultaneously against different competitors in order to avoid the collapse of the system. This kind of circular dependence requires measuring all parameters for all species to be able to estimate its behaviour. Even when these parameters are correctly measured spending long hours in the field, the predictive power of such mechanistic models is still very low (See S1 Appendix at Supplementary Material).

Alternatively, it is possible to use data-driven predictive models where the response variable (species abundance in our case) could also be a function of abiotic and biotic features. While this distinction among features is ecologically important in terms of the ultimate mechanisms driving species abundances, from the point of view of the data scientist that distinction is not relevant, as far as the model behaves properly. In our particular scenario, the hypothesis is that the abundance of any given plant species is influenced by the environment (e.g. precipitation, soil properties) and the abundance of competitors. In mathematical terms:

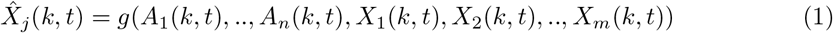

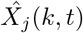 is the predicted abundance of species *j* in the subplot (our spatial sampling unit) *k* at instant *t*, and *g* is the predictor, a function with *n* abiotic variables and *m* abundances of competitors, including individuals of *X_j_*(*k, t*). If the available dataset includes all these variables, the data engineer enjoys a wide range of chances to pick out of them a subset of features to train and tune the model. However, while abiotic variables are often easy to measure in a spatially explicit way, obtaining spatially explicit data on the species abundance of the whole community is prohibitive, and in fact, it would be equivalent to measure community composition to predict community composition. For that task, a predictive model would be useless.

In any case, and for the sake of being pedagogic, we start by testing the scenario where the full dataset is available, and the field team recorded a detailed sample of species abundance and abiotic parameters for each area of interest. In this case it is simple to build a predictive model that works for a nearby piece of land, where all those variables are known: this is the very essence of machine learning. So, abundance at *t* of species *j* in a given area *m* is estimated by:

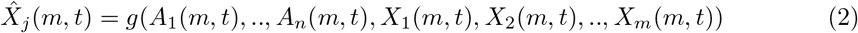

But as stated above, things get harder when trying to apply the model to common prediction challenges outside of the sampled area. For example, how do we know in advance the abundances of competitor individuals elsewhere in the community, or the abundance of the plants that will ripe next year?

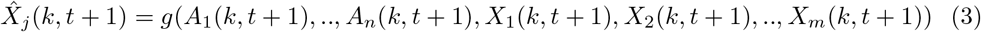

Equation 3 is deceptively simple, because none of the variables are known at *t* + 1 (next year). These data are unavailable by definition, we need to predict them, so there are *N* +1 prediction problems instead of 1. Abiotic features may be gathered next season without an extraordinary effort, but if we need to know the exact values of the abundance of competitors *X*_1_(*k, t* + 1), *X*_2_(*k, t* + 1), …, prediction is pointless.

To put it bluntly, imagine you have sampled 100 areas (i.e.subplots) out of 10000 to build a detailed map of density of one species that has 20 competitors. The sample is representative of the population and there are no quality issues. Even with that optimal starting point, you need to count the individuals of competitors for each of the unknown 9900 plots. The only way to avoid that time-consuming task is predicting those abundances, but then you face 23 unsolvable problems.

A possible strategy to overcome the deadlock is dropping off the conflicting variables. That is, getting rid of the species abundances and relying just on abiotic data.

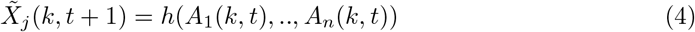

From an ecological perspective this solution may seem too radical, as the model ignores species interactions. For the data scientist, feature engineering is a common procedure to build and test different models. Data sets have redundant information and dimensionality reduction is often desirable. Therefore, we can start building a model with only abiotic predictors for spatial application. Even from an extreme data-centric approach, this solution looks very weak for this predictive challenge. But *weak* doesn’t mean *useless*. A smart mix of weak models may produce an accurate predictor, that is the basis of ensemble methods [33]. This first model generates a set of abundances driven only by abiotic factors. In a second step, we predict again species abundances with the same abiotic data and the predicted abundance of competitors modeled in step one. Thus, we end up with a two-step predictor that is an *ad-hoc* ensemble method for this scenario. To summarize, first we predict abundances of competitors in the unsurveilled subplot *p* using a model built just with abiotic data as in equation 4, and second, we combine the observed abiotic data with the abundances predicted from the first model:

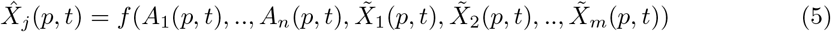

Here we advance some key results showing that the two-step predictor is much more accurate than the one based on abiotic data. Its global performance is close to that of the global predictor fed with observed quantities, not mere weak predictions.

## Materials and methods

### Prediction

We tracked during five years (2015-2019) the local abundances of individuals of 23 annual plant species distributed along 9 plots (plot size 8.5m x 8.5 m) in a highly diverse Mediterranean grassland located along a salinity gradient of 1km long by 800 m wide. Each plot is subsequently subdivided in 36 subplots of 1 *m*^2^. For each of these subplots, we compiled across each of these five years the number of adult individuals of each plant species at their phenological peak (i.e. when at least half of the individuals are in bloom). This period extends on average and across species from February to June yearly. In addition, we empirically measured an array of physical and chemical soil properties at the beginning of the survey (spring 2015) to characterize the abiotic properties of each plot (see Table S2), and we finally obtained annual precipitation values for all years from a nearby weather station maintained by the regional government “Junta de Andalucía” (El Rocío-Almonte, 10 km far apart).

In total, the dataset contains abundance values for 37240 species-plot combinations. The distribution of abundances is extremely skewed due to a 75.6% of zero values, meaning that most species are very scarcely represented because they were only recorded some years and in some particular plots of the soil salinity gradient. This is a well-known problem in spatial distribution models [34]. Removing them, the uneven distribution of abundances remain, as generally expected from species-abundance distributions (Fig. 1A). As expected, the mean value and the variance of abundances scale with each other. This phenomenon is known as Taylor’s Law and in our case, the scaling an exponent of 2.15 and an adjusted *R*^2^ = 0.92 [35]. Taylor’s Law appears in different contexts in ecology with exponents close to 2 as in this case [36, 37], which implies that our sampling is representative of empirical community structures.

**Fig. 1.**
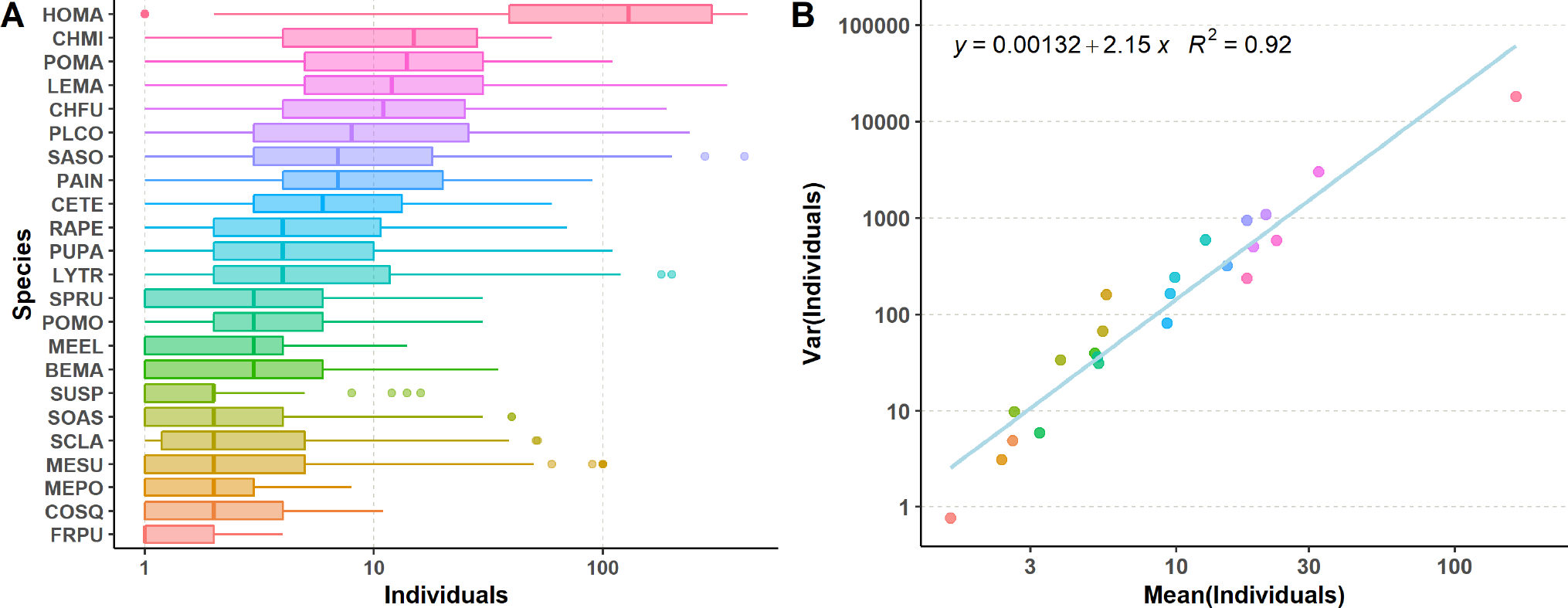
Species abundances. A: Boxplots of the individuals distribution for each species, highlighting the median value. B: Scatter plot of the mean vs. variance for individuals by species, and regression line to check how they fit Taylor’s Law.

### Methods

#### Regression models

We implemented three regression models to tackle the problem to predict species abundances in space and time. Linear Regression, Random Forest regression and XGBoost use the equation presented in 5. Specifically, linear Regression Model (LRM) is a good choice to achieve a balance between interpretability and precision. It explains the outcome as a function of the multiple input features and has inspired many mechanistic models. This simple model provides fair results when the underlying function is linear or there are linear combinations of features.

Therefore we also used more flexible models to improve results. Random Forest Regression (RFR) is a tree-based ensemble method, and belongs to the family of Classification and Regression Trees (CART) [38]. It generally works fine with high-dimensional problems. It combines the predictions from multiple weak trees to make accurate predictions [39]. A random subset of samples is drawn with replacement from the training sample. All of them have the same distribution. These randomly selected samples grow decision trees and the average of predictions yields the model’s outcome [40]. Alternatively, XGBoost (eXtreme Gradient Boosting) relies on the concept of gradient tree boosting [41, 42]. Boosting is a sequential algorithm that makes predictions for T rounds on the entire training sample and iteratively improves the performance of the boosting algorithm with the information from the prior round’s prediction accuracy. It is faster to train and less prone to overfitting than a Boosted Regression Tree (BRT) [43]. XGBoost produces black models, hard to visualize and tune compared to RFR.

One common feature to all these methods is that they are sensible to the random splitting of training and testing sets, which we typically set to 80/20 ratio. Then, for each model we perform a 7-fold cross validation [44]. In addition, we provide the results of 100 of such models (i.e. hereafter runs) and not of just one run.

#### Feature engineering

The dataset which fed this regression analyses includes 40 variables. There are 14 abiotic measurements, 13 of soil conditions (pH, total salinity, carbonates, Organic mater, C/N ratio and Cl, C, N, P, Ca, Mg, K, and Na concentrations; Table S2) for each subplot, and the annual rainfall, common for all plots. The additional 23 numerical features are the abundances of each species in the subplot (Supplementary table S3).

For the three models (Linear Regression, Random Forests and XGBoost), we tried to implement variable or feature selection methods. Those methods reduce the model complexity dropping those variables that are less relevant for the outcome. Among the most widely used are the wrapper model [45], the filter model [46] and the hybrid model [47]. The wrapper model uses the predictive accuracy of a predetermined learning algorithm to determine the goodness of the selected subsets. The filter model separates feature selection from classifier learning and selects feature subsets that are independent of any learning algorithm. Hybrid algorithms focus on combining filter and wrapper algorithms. We use filtering for its computational efficiency against the wrapper and hybrid models. The analysis of Spearman’s Correlation showed up that there were no significant dependencies among this set of variables (Figs. S1, S2). Permutation Importance technique tests the performance of a model after removing each feature and replacing it with noise [48]. Around 25% of variables provided more information than random noise, both for the abiotic model Table 1, Supplementary table S4) and the full model. Annual rainfall proved to be one of the most relevant features, as well as the species or the presence of other individuals. Certain species such as POMA, LEMA, CHFU and SASO (see Supplementary table S3 for species acronyms) or the concentration of carbonates showed up to be relevant too. Although feature analysis helped to identify which of them where more relevant, feature reduction did not improve predictive models for this particular dataset. So, we included the aforementioned 14 abiotic and 23 biotic features in the final models.

**Table 1.** Feature importance for the ABIOTIC model

| Feature | Importance |
| --- | --- |
| *Species* | 0.328 |
| *Annual precipitation* | 0.251 |
| *Carbonates* | 0.086 |
| *P* | 0.054 |
| *Organic matter* | 0.039 |
| *Random noise* | 0.032 |
| *Salinity* | 0.030 |
| *C/N ratio* | 0.029 |
| *K* | 0.028 |
| *C* | 0.023 |
| *Ca* | 0.021 |
| *Cl* | 0.021 |
| *Na* | 0.019 |
| *Mg* | 0.018 |
| *pH* | 0.016 |
| *N* | 0.005 |

### Model evaluation

To assess the performance of regression models we compute the Root Mean Square Error (RMSE) and the Relative Squared Error (RSE) [49]. RMSE is a distance between the vectors of predicted values (*y_i_*) and observed values 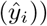.

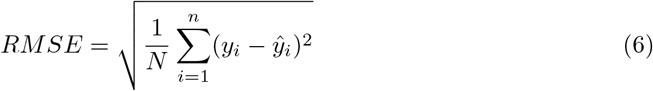

The RSE normalizes RMSE by dividing it by the total squared error of the predictor. It is more convenient to compare models, when the outcome is so skewed. Moreover, it enables comparisons among species whose abundances are rather heterogeneous.

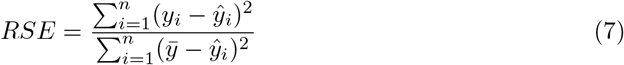

#### The two-step model

As we mentioned above, prediction of abundances in this scenario poses a major challenge as the problem is recursive. To predict the abundance of species *X* we need to know in advance the abundance of each of its competitors, but those abundances are dependent on the rest of species as well.

To solve this limitation and given the fact that soil features and annual rainfall are easier to get, a predictor that could get rid of all abundances is more operative, paying the price of a reduced quality. Dropping that information is equivalent to ignore direct interactions among species. That would be unacceptable for a mechanistic model as a too naïve simplification, but Machine Learning has developed some strategies to deal with this kind of hindrances. Stacked models are a kind of ensemble models that perform sequential learning [50]. Predicted values of stage *n* are fed as features to stage *n* + 1 mixed with original features. We have built a two-step sequential model, following this idea. During the first step, a Random Forest is trained with the abiotic data, and predicts the abundances of competitor individuals. These predictions are weak, but combining them with the abiotic variables, the semi-synthetic dataset trains an all-features model to perform the final prediction, using Linear Regression, Random Forest or XGBoost.

The *two-step model* is evaluated as the other methods using RMSE and RSE, but we needed to introduce a refinement to measure its performance for each species. As there are a 75% of zero values of the *individuals* feature, the abiotic first step models tends to predict zero abundances for rare species. To avoid this, we introduce resampling SMOTE correction procedure [51]. This solution leads to a better generalization during the first stage prediction. The SMOTE correction is just used for this second purpose, the general two-step model do not over sample in any way.

In the final step of the analyses, we build full predictors to evaluate spatial prediction by randomly splitting the data in training and testing data using the cross-validation explained above. For temporal prediction, the training set excludes the samples of the year we want to predict.

## Results

We found that models perform quite well regarding their accuracy to predict species abundances within a spatial context. This is an important result because it shows a direct application of using Machine Learning approaches to describe relevant characteristic of ecological communities such as the spatial distribution of species relative abundances. Specifically, we build 100 models that only differ in the random split of training and testing sets, including all features and years. The median *RSE* values are 0.898 for the Linear Regressor, 0.130 for XGBoost and 0.134 for Random Forest (table 2). This implies that under the best model, we can predict plant species abundances with an error of 13%. Prediction would be a practical tool with these two last models, but results may be deceiving to ecologists. To predict the abundance of species *X* we need to know beforehand the abundances of the rest of species, so the painstaking field work is not avoided.

**Table 2.**
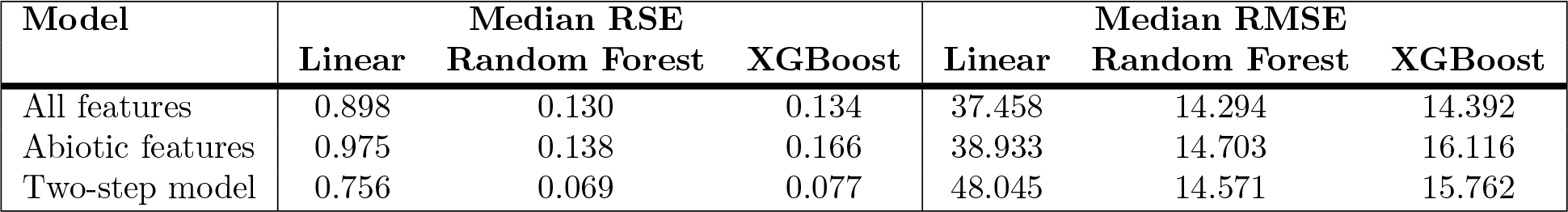
Prediction errors for spatial application

The median RSE value for the Random Forest predictor trained just with abiotic information is roughly the same of the predictor with all features: 0.138 vs. 0. 130. This figure could be misleading because of the skewed distribution of abundances, but provided the hint to try the two-step method. This stacked generalization predicts the abundances of competitor species using the abiotic Random Forest model. Results are quite encouraging as median RSE falls to 0.069 using Random Forest for the second stage and to 0.077 using XGBoost. The practical advantage of the two-step method is that it does not require to know in advance the abundance of competitor species.

Fig. 2 shows the improvement of the RSE distribution with the two-step method and Random Forest as the second stage model. XGboost results were slightly worse (Supplementary Figs. S3, S4).

**Fig. 2.**
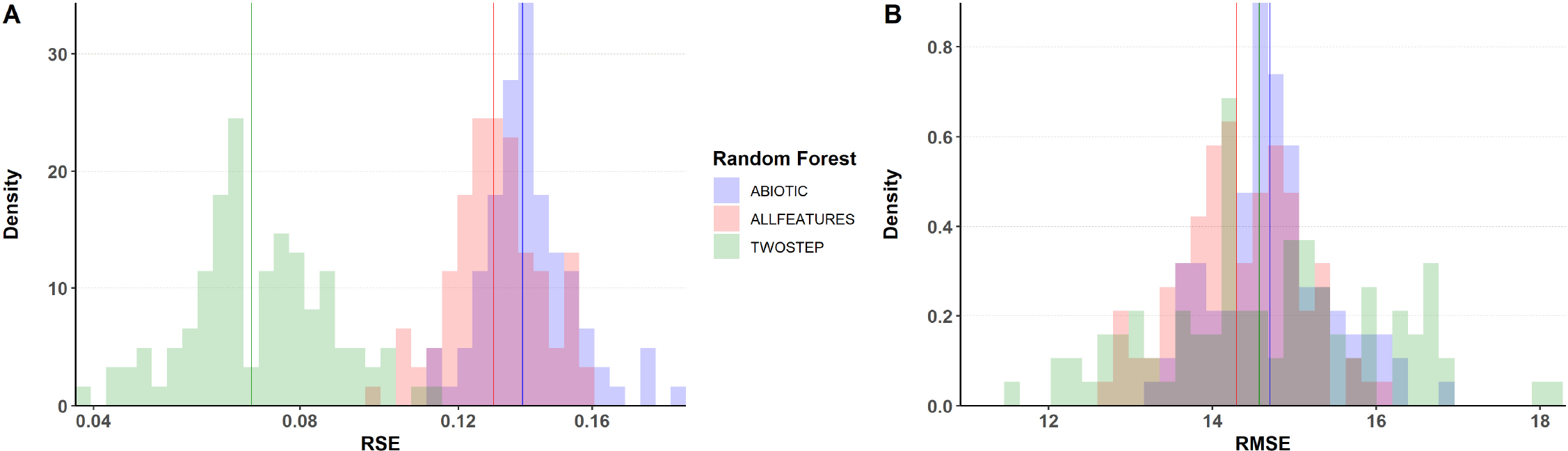
Prediction errors with a two-step Random Forest Regressor. A: Relative Squared Error distributions for 100 random choices of training/testing sets, vertical lines set at median values. B: Root Mean Square Error distributions for the same collection of predictors.

Although RSE is useful to make global comparisons among predictors (i.e. among species), we still require an assessment of prediction accuracy by species because of their asymmetry in observed abundances. To evaluate the three methods considering a species-specific approach, we performed 100 runs, following the steps described in the previous section, and measured both RMSE and RSE for each species. We overall found that relative prediction error is fairly small for abundant species such as *Hordeum marinum* or *Chamaemelum fuscatum*, while it shows a strong spread for plants that are relatively rare in the study area (Fig. 3, see also Supplementary Figs. S5, S6).

**Fig. 3.**
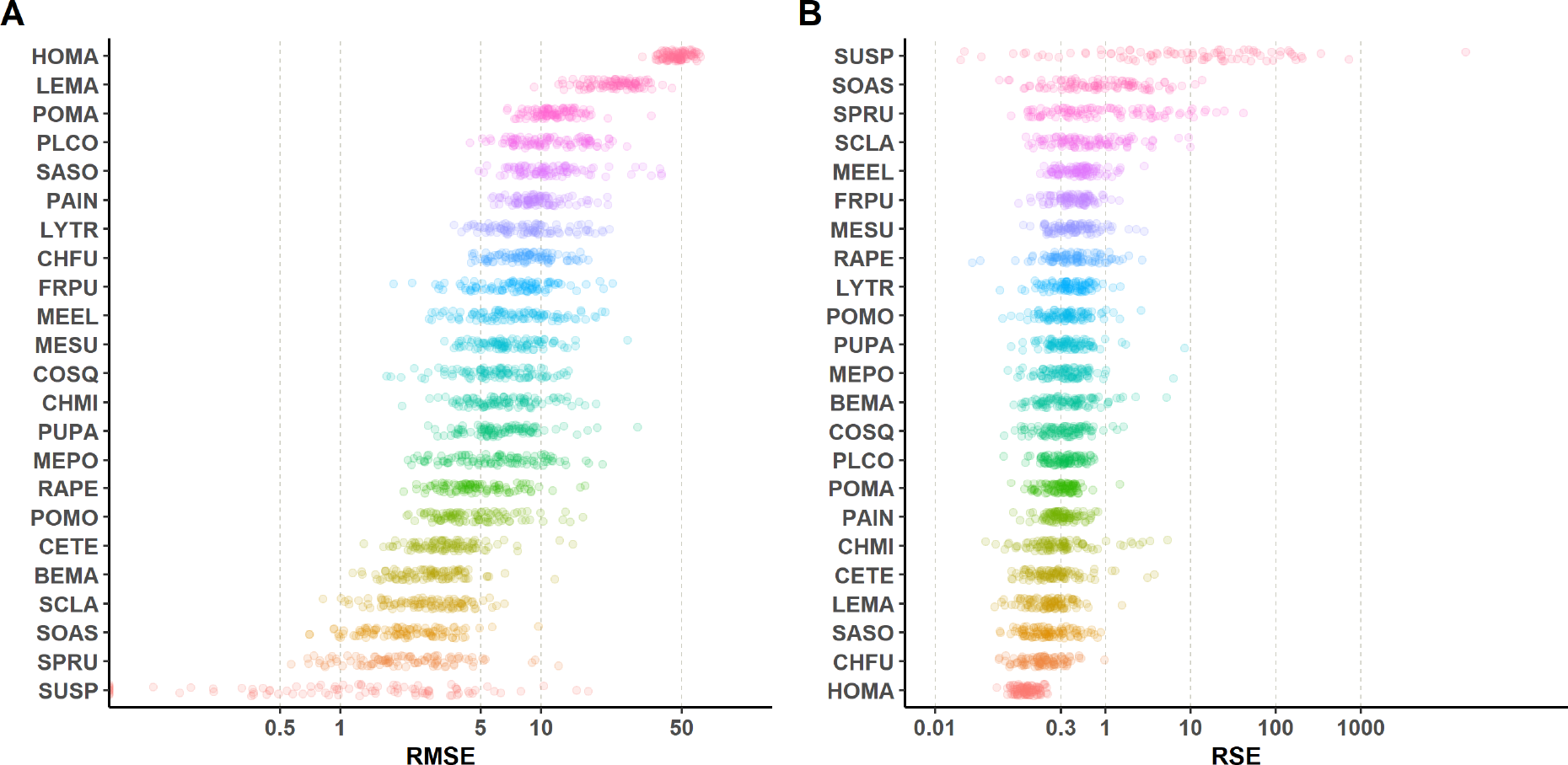
Prediction errors by species using a two-step Random Forest regressor. A: Relative Squared Error distributions for 100 random choices of training/testing sets. B: Root Mean Square Error distributions for the same collection of predictors.

Similarly to predicting species abundances across space, we aim to predict species abundance over time. From a modelling perspective, prediction over time is a widespread application of Machine Learning but also one of the toughest. One key condition is that the underlying system must show some kind of seasonal behaviour superimposed to a long-term trend. Our ecological study system fortunately meets this requirement because both abiotic and biotic variables follow an annual cycle, which is reset every year with the fall rains. Therefore, if we have got a curated yearly series of data, it is straightforward to build a predictor for the incoming season, and in the case the quality of predictions is fine enough, then it would allow us to anticipate how plant will respond to changes in future environmental conditions.

Unfortunately, this expectation is not the case for the data analyzed, and it comes as no surprise. Table 3 shows the evaluation results of temporal predictors, trained with all features of four years and tested with the remaining one. If we include the Precipitation variable, the RSE using Random Forest regression ranges from a disappointing 0.60 to a staggering 6.30. This may be interpreted as the best prediction is off by roughly a 60%. A potential explanation is that, despite the fair size of the dataset, the temporal sample is tiny. In addition, yearly fluctuations in weather are heavily marked in this study system, ranging from 384 mm in 2019 to 625 mm in 2016. The fact that there are only five values for time-related variables, one per year, makes prediction to fail because the test data often falls outside the trained data conditions. One possible workaround is dropping this variable to reduce overfitting. Results show a mild improvement but even the best RSE (0.44 for 2016) is still too weak. Results for Linear Regression and XGBoost predictors are even worse.

**Table 3.** Prediction errors splitting by year and using Random Forest

| Predicted Year | With Precipitation |  | Without Precipitation |  |
| --- | --- | --- | --- | --- |
|  | RMSE | RSE | RMSE | RSE |
| 2015 | 37.67 | 6.30 | 35.73 | 5.67 |
| 2016 | 31.51 | 4.99 | 27.17 | 3.71 |
| 2017 | 38.25 | 0.60 | 32.99 | 0.44 |
| 2018 | 34.29 | 0.75 | 34.92 | 0.77 |
| 2019 | 44.70 | 0.61 | 39.45 | 0.48 |

Regardless of the differences in the ability of the Random Forest models to predict species abundance over time or across space, these models have the potential to provide novel insights into some key processes that modulate the response variable studied (species abundances in our case). This new information can be incorporated in turn into mechanistic predictions from population dynamics models that describe the abundance trajectories of interacting species. These later type of models are much more familiar to ecologists. This possibility of feedback from the data-driven models to the mechanistic models is exemplified in our system with the particular focus on soil carbonates. The inclusion of this abiotic variable, which was deemed important by the data-driven model (Table 1), shows an overall improvement in the temporal predictions derived from the mechanistic models (S1 Appendix). Hence, we advocate for a bidirectional learning process between data-scientists and ecologists.

## Discussion

By combining ecological knowledge with data-driven models, we showed that it is possible to develop simple models that predict well complex systems such as the abundance of multiple species that compose ecological communities. Plant species composition at fine-resolution scales is hard to predict, because their densities and relative abundances are partly governed both by biotic factors, which determine where species can potentially thrive, and by the network of species interactions in which they are embedded, which modify their reproductive success. In fact, these two axes of variation defining the species persistence probabilities have been at the core of the species niche concept [52], and in the development of modern community ecology theory [53], but rarely exploited for predictive purposes. Here, we show a simple methodology to use easy to obtain abiotic information to accurately predict species abundances while taking also into account their potential biotic interactions. Our models are surprisingly accurate for spatial predictions with less than a 10% error in predicting species abundances, but limited for temporal predictions because of the short span of the dataset.

Machine learning-based methods have been extensively applied for relating species distributions to environmental factors, through species distribution models. While the literature on species distribution modelling is vast, most of it is centered on large scale distributional patterns of species occurrences [54], often involving only abiotic variables [55], and in a vast majority of cases, prediction is limited to species presence or absence. However, most ecosystem functioning processes happens at the community scale. At this scale, species interactions are thought to determine species performance, quantified in their probability of persistence [27] and in their relative abundance [56]. We show that a data-driven sequential model that firstly predicts the potential species abundances for a given set of abiotic variables, and secondly uses this predictions to refine the realized species abundances predicted, performs incredibly well and even better than more data-hungry models. The fact that this two-step process enhances predictions over a one-step model with all data available is remarkable. One possible explanation is that observed plant abundances empirically measured in the field only capture fully developed individuals, missing early stages of competition among seedlings that despite dying soon, affect final species abundances. Similarly, we use the number of individual competitors per species but we did not account for their size, something that normally occurs when considering species such as trees that show a greater range in individual sizes compared to annual plants. The two-step process may better indirectly capture the size variation during the modelling process.

As expected, the best performing data-driven models are Random Forests (followed closely by XGboost). The assumptions of linear models are too simple when there are complex interactions among features as the exploratory analysis suggested. Interestingly, this data-driven exercise can also help us enhance mechanistic models. We already used mechanistic models to understand the species dynamics in our ecological system. Aware of the importance of the abiotic environment, we modelled species reproductive success as a function not only of competitors, but also of other environmental variables such as soil salinity content [16]. To our surprise, the feature importance selection procedure highlights *CaCO3* as a key determinant of species abundances and not salinity, which was the most obvious variable initially selected in the field. Despite initially counter-intuitive, this result is congruent wit the fact that we sampled in a hypersaline environment in which phosphorous (a key element for plant growth) is not available for plant absorption. Rather, it is retained in carbonate minerals such as calcite and dolomite, and plants can mostly obtain phosphorous thanks to the enzymes from mycorrhizal fungi. With this new knowledge, we re-parameterized the mechanistic annual plant model by adding *CaCO*3 as a covariable affecting both the intrinsic fecundity rates and the pairwise interactions among species. With this update we obtained significantly better predictive error than with the biotic-only parameterization (Table S1). Hence, we show that ecological process can inform the data driven models, but those can in turn refine which ecological process are important to include int the mechanistic models.

This exercise is tailored to the problem at hand. For example, an implicit assumption of this modelling framework is that plant species can reach all quadrants in the grassland, and are not limited by dispersal. This assumption is reasonable on a study system in which seeds are small, they can be dispersed by wind and small animals such as ants, and additionally the system also gets flooded in extremely wet years. Similarly, we focused our modelling on the plant-plant competitive interactions, which are the main interactions structuring this grassland communities [57]. However, the same approach can be used to model other interaction types in other systems, as far as you have initial data to train the models. In our case, we obtain a good spatial predictive ability, but we fail to predict temporally. Given the strong across-year variations in precipitation, we believe this is due to the limited number of years to train the data, and not an inherent limitation of the framework. Hence, once we compile longer temporal series, we are confident it will be likely to obtain models that accurately predict temporal variation in species abundances. It might also be possible that stochastic events, which create variation from unknown sources (e.g. random birth-death, perturbations in population sizes, dispersal events in no particular direction) are more prevalent in the temporal dimension than deterministic processes such as species interactions. In any case, given the expected ongoing environmental change in many abiotic variables such as precipitation regimes and temperatures, the prediction ability of these models will be crucial to anticipate to global change effects on delicate and highly-diverse ecosystems such as Mediterranean grasslands.

## Conclusion

The rate of ecological data generated is increasing substantially [58]. Open and reliable datasets hold the potential to facilitate the application of near-term forecasting protocols [59]. However, for those efforts to thrive, we need simple models that can work with the sparse data typical of ecological surveys. A more predictive ecology likely serves to anticipate how several ongoing critical environmental changes such as climate change affect multiple properties of ecosystems, and at the same time it also provides information about which management actions are required to maintain healthy ecosystems. Taken together, our results show that two-step ensemble models are a promising tool to reach efficient management without the costs of prohibiting data collection.

## Supporting information

**S1 Fig. Abiotic features correlation.**

**S2 Fig. All features correlation.**

**S3 Fig. Prediction errors with a Linear regressor.**

**S4 Fig. Prediction errors with a XGBoost regressor.**

**S5 Fig. Prediction errors by species using a Linear regressor.**

**S6 Fig. Prediction errors by species using a XGBoost regressor.**

**S1 Table. Prediction errors for annual plant models with and without the inclusion of CO3 as an abiotic driver.**

**S2 Table. Abiotic variables**

**S3 Table. Plant species**

**S4 Table. Feature importance for the Random Forest predictor with all variables.**

## Authors’ contributions

IB, OG and JG-A conceived the idea, OG and IB collected the data. IC, JG-A, DG-C and JG analysed the data. IC developed the predictive models and JG-A the visualizations. JG-A and IB lead the writing with inputs of all co-authors.

## Acknowledgments

O.G. acknowledges support provided by the Ramón y Cajal Programme (RYC-2017-23666). I.B., O.G. and J.G-A also acknowledge financial support provided by the Spanish Ministry of Economy and Competitiveness Explora Program (CGL2017-92436-EXP, SIMPLEX) and Retos Investigación (RTI2018-098888-A-I00, MeDiNaS). J.G. acknowledge financial support provided by the Spanish Ministry of Science, Innovation and Universities (PGC2018-093854-B-I00). We thank Doñana National Park staff for granting access to Caracoles real estate.

## Data availability and code

Code and data are available at https://doi.org/10.5281/zenodo.4609824 Reproducibility instructions are detailed in the README.md file.

